# A neuromorphic model of olfactory processing and sparse coding in the Drosophila larva brain

**DOI:** 10.1101/2021.06.29.450278

**Authors:** Anna-Maria Jürgensen, Afshin Khalili, Elisabetta Chicca, Giacomo Indiveri, Martin Paul Nawrot

## Abstract

Animal nervous systems are highly efficient in processing sensory input. The neuromorphic computing paradigm aims at the hardware implementation of neural network computations to support novel solutions for building brain-inspired computing systems. Here, we take inspiration from sensory processing in the nervous system of the fruit fly larva. With its strongly limited computational resources of <200 neurons and <1.000 synapses the larval olfactory pathway employs fundamental computations to transform broadly tuned receptor input at the periphery into an energy efficient sparse code in the central brain. We show how this approach allows us to achieve sparse coding and increased separability of stimulus patterns in a spiking neural network, validated with both software simulation and hardware emulation on mixed-signal real-time neuromorphic hardware. We verify that feedback inhibition is the central motif to support sparseness in the spatial domain, across the neuron population, while the combination of spike frequency adaptation and feedback inhibition determines sparseness in the temporal domain. Our experiments demonstrate that such small-sized, biologically realistic neural networks, efficiently implemented on neuromorphic hardware, can achieve parallel processing and efficient encoding of sensory input at full temporal resolution.

## Introduction

Neuromorphic computing [1] is a novel paradigm that aims at emulating the naturalistic, flexible structure of animal brains on an analogous physical substrate with the potential to outperform von Neumann architectures in a range of real-world tasks [2, 3]. It can inspire novel AI solutions [4–6] and may support control of autonomous agents by spiking neural networks [7–9]. A major challenge for brain-inspired neuromorphic solutions is the identification of computational principles and circuit motifs in animal nervous systems that can be utilized on neuromorphic hardware to exploit its benefits.

Drawing inspiration from neural computation in the nervous systems of insects is particularly promising for developing neuromorphic computing paradigms. With their comparatively small brains ranging from ≈ 10,000 neurons in the fruit fly larva to ≈ 1 million neurons in the honeybee, insects are able to solve many formidable tasks such as the efficient recognition of relevant objects in a complex environment [10, 11], perceptual decision making [12–14], or the exploration of unknown terrain and navigation [15–19]. They also show simple cognitive abilities such as learning, or counting of objects [20–24]. At the same time, their compact nervous systems are optimized for energy efficient computation with limited numbers of neurons and synapses, making them ideally suited to meet current neuromorphic hardware limitations regarding network size and topology. Spiking neural networks modelled after the insect brain have been shown to support efficient sensory processing [25], learning [7, 26], foraging and navigation [27–29], and counting [28]. Model studies also include earlier neuromorphic implementations of insect-inspired computation [4, 5, 9, 30–33].

Sparse coding [34, 35] is a fundamental principle of sensory processing, both in invertebrates [36–40] and vertebrates [41–45]. By transforming dense stimulus encoding at the receptor periphery into sparse representations in central brain areas, the sensory systems of animals achieve energy efficient and reliable stimulus encoding [35, 46], which increases separability of items [47–50]. Sparse coding in neural systems has two major components [39]. *Population sparseness* refers to the representation of a stimulus across the entire population of neurons, such that only few neurons are activated by any specific stimulus and different stimuli activate largely distinct sets of neurons. Re-coding from a dense peripheral input to a sparse code in central brain areas supports stimulus discriminability and associative memory formation by projecting stimulus features into a higher dimensional space [51–53]. *Temporal sparseness* indicates that an individual neuron responds with only a few spikes to a specific stimulus configuration [34, 54, 55] supporting the encoding of dynamic changes in the sensory environment [42, 56] and memory recall in dynamic input scenarios [28].

We are interested in the transformation of a densely coded input into a sparse representation within an olfactory pathway model of the *Drosophila* larva. As a common feature across insect species, odor information is processed across multiple network stages to generate a reliable sparse code of odor identity in the mushroom body [36, 57, 58], a central brain structure serving as a hub for multi-sensory integration, memory formation and memory recall [10, 59]. A shared characteristic of the *Drosphila* larva brain and the here-used real-time neuromorphic hardware system is their relatively small network size. With this limited capacity, computational efficiency and frugal use of the limited resources are a major constraint. Implementing evolutionary-derived mechanisms from the insect brain that allow for sparse, thus more efficient stimulus encoding on the chip could help to broaden the scope of its applications. In our network model we test the efficiency of cellular mechanisms and network motifs in producing population and temporal sparseness and test their implementation on the mixed-signal neuromorphic hardware DYNAP-SE [60] in comparison to a software simulation using the Python-based spiking neural network simulator “Brian2” [61].

## Methods

### Spiking neural network model

The architecture of the spiking neural network model as shown in Fig. 1 A uses the exact numbers of neurons in each population and the reconstructed connectivity for one hemisphere as published in the electron-microscopic study of a single animal [62]. The network consists of 21 olfactory receptor neurons (ORN) at the periphery, 21 projection neurons and 21 local interneurons (LN) in the antennal lobe and 72 Kenyon cells (KC). In each brain hemisphere there is exactly one anterior paired lateral neuron (APL). We hypothesize that the APL receives input from most or all mature KCs [63] included in this network model. Due to technical limitations of the DYNAP-SE chip with a maximum in-degree of 64 synapses for one neuron we randomly chose 64 KCs that provide input to the APL. This choice was fixed for the model, both on the hardware network and in the software simulation. We further hypothesize, based on evidence in the adult species, that all ORNs and all KCs have a mechanism of cellular spike frequency adaptation (SFA).

**Fig. 1.**
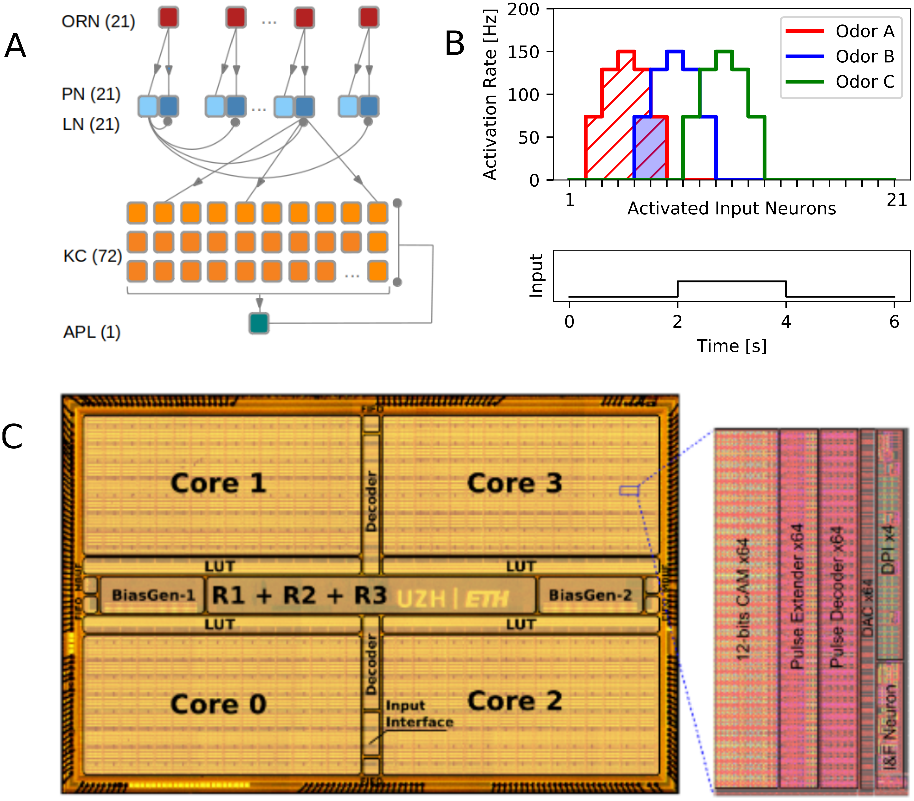
Neuromorphic spiking neural network approach. (A) Network model of the *Drosophila larva* olfactory pathway including all neurons and connections implemented. One-to-one feed-forward connections between olfactory receptor neurons (ORN, red) and projection neurons (PN, dark blue)/local interneurons (LN, light blue) and from PN to Kenyon cells (KC). Lateral inhibition from each LN to all PN and feedback inhibition from the anterior paired lateral neuron (APL) to KCs. The number of neurons in each population is declared in parenthesis. (B) Input pattern of the three artificial odors used and time course of the odor stimulation protocol (excluding the warmup) with odor onset at 2s and offset at 4s (lower panel). The odors are characterized by their ORN activation profile and implemented with varying degree of similarity (overlap as indicated by the shaded area).(C) Chip micro-photograph of the DYNAP-SE device. The chip, fabricated using a standard 180 nm CMOS technology, comprises four cores with 256 adaptive exponential integrate-and-fire neurons each. The inset shows a zoom into an individual neuron with an analog neuron circuit, analog synapse circuits and digital memory and communication blocks. The central part of the chip contains the asynchronous routers for transmitting spikes between individual neurons and bias generators with 12 bit current mode DACs for setting the network parameters.

### Implementation on the DYNAP-SE neuromorphic hardware

The olfactory pathway model of the *Drosophila* larva was implemented using the Dynamic Neuromorphic Asynchronous Processor (DYNAP-SE) [60] (Fig. 1C). This processor is a full-custom mixed-signal analog/digital VLSI chip, which comprises analog circuits that emulate neurons and synapses with biologically plausible neural dynamics. Given the analog nature of the circuits used, the synapses and neurons exhibit parameter variability that is characteristic also of real neurons. The analog circuits used, implement multiple aspects of neural dynamics, such as spike-frequency adaptation (implemented as a shunting inhibitory synapse), refractory periods, exponentially decaying currents, voltage-gated excitation and shunting inhibition [60, 64]. The silicon neurons circuits, similar to their biological counterparts, produce spikes. In the chip, these are stereotyped digital events which are routed to target synapses by a dedicated Address Event Representation (AER) infrastructure [65, 66]. The conductance-based synapses are current-mode circuits [64] that produce an EPSC with biologically plausible dynamics, which are then injected into the neurons *leak* compartment. This compartment acts as a conductance block which decreases the input current as the membrane potential *v_i_* increases. One of the inhibitory synapses subtracts charge directly from the membrane capacitance and provides a shunting inhibition mechanism [64]. All other synaptic currents are in turn summed together and integrated in the post-synaptic neurons leak compartment.

The model (Fig. 1 A) was initially developed in software and the neural architecture was then mapped onto the mixed-signal hardware by configuring the AER routers and programming the chip digital memories to connect the silicon neurons via their corresponding synapses. The parameters of the hardware setup were fine-tuned using the on-chip bias generator, starting from the estimates provided by the software simulation. Computer-generated control stimuli, in the form of well defined spike trains, were provided to the chip via a custom FPGA (Field Programmable Gate Array) board. Each neuron population was implemented on a single core, using in total five cores and two chips. All the circuit biases of neurons belonging to different cores could be tuned independently. The synapses from ORN to PN, PN to KC, and KC to APL were designed as excitatory whereas the synapse from LN to PN and APL to KC were implemented as inhibitory. SFA was implemented in the ORN and KC neuron populations.

Three separate recordings were done, one for each of the three odors (Fig. 1 B). Within each of the three experiments all six conditions (different sparseness mechanisms enabled) were recorded always in the same order (LN+APL+SFA, LN+SFA, LN, APL+SFA, LN+APL, SFA).

### Computer simulation of the spiking neural network

The simulations were implemented in the network simulator Brian2 [61] and run on a X86 architecture on an Ubuntu 16.04.2 Server. All neurons (Fig. 1 A) were modeled as leaky integrate-and-fire neurons with dynamic synapses. The membrane potential *v*^i^ obeys a fire-and-reset rule, being reset to the resting potential whenever the spike threshold is reached. The reset is followed by an absolute refractory period of 2 ms, during which the neuron does not integrate inputs (Table 1). The membrane potential of a neuron in a particular neuron population (v^O^, v^L^, v^P^, v^K^, v^A^) is governed by the respective equation. The neuron parameters can be found in Table 1.

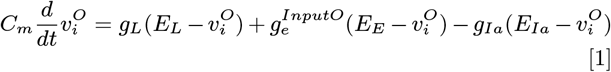

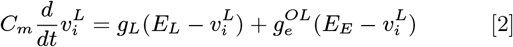

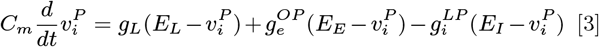

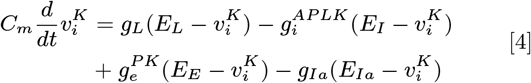

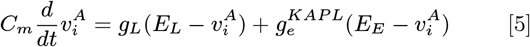

where ORNs (Eqn. 1) and KCs (Eqn. 4) are equipped with an additional spike-triggered adaptation (Eqn. 6) where *g_Ia_* is the adaptation conductance and *τ_Ia_* is the decay time constant. With every spike *g_Ia_* is increased in ORNs and KCs by 0.1 nS and 0.05 nS respectively.

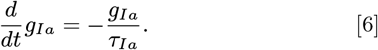

**Table 1.**
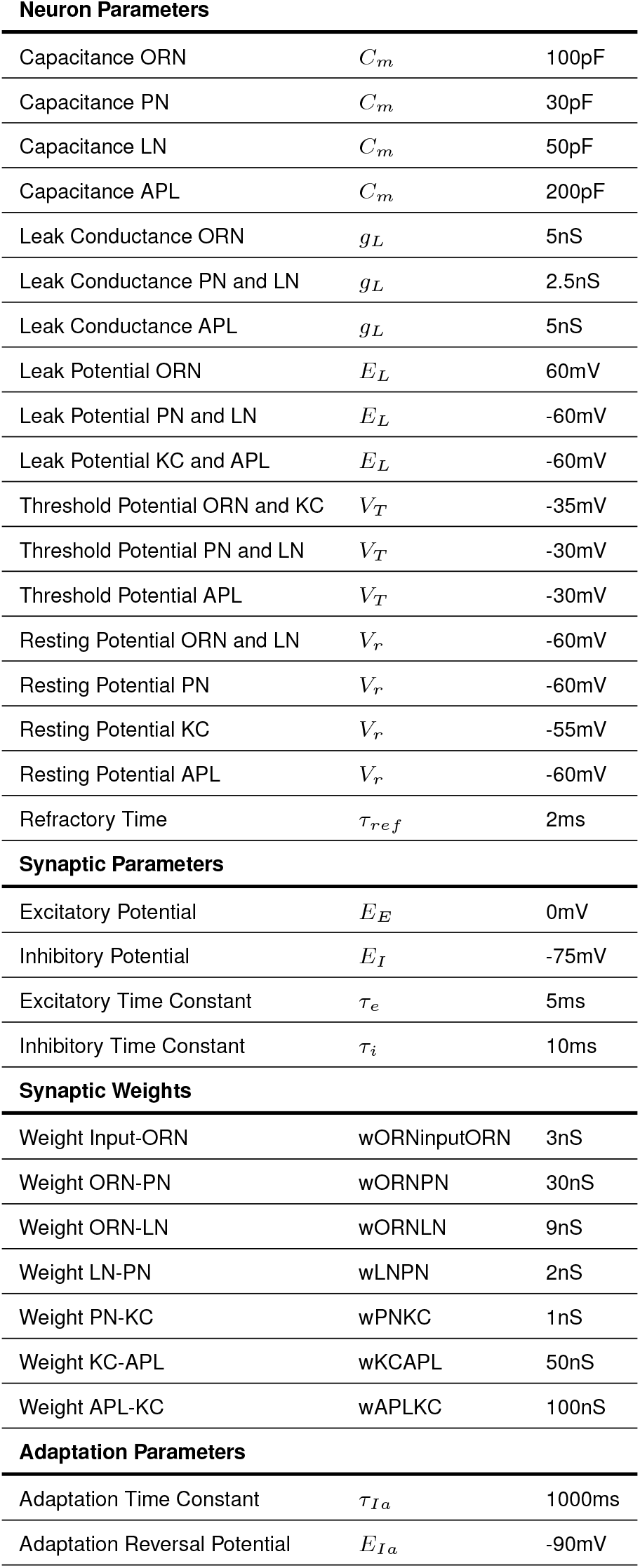
Network simulation parameters.

Neuron Parameters
|  |  |  |
| --- | --- | --- |
| Capacitance ORN | $C_m$ | 100pF |
| Capacitance PN | $C_m$ | 30pF |
| Capacitance LN | $C_m$ | 50pF |
| Capacitance APL | $C_m$ | 200pF |
| Leak Conductance ORN | $g_L$ | 5nS |
| Leak Conductance PN and LN | $g_L$ | 2.5nS |
| Leak Conductance APL | $g_L$ | 5nS |
| Leak Potential ORN | $E_L$ | 60mV |
| Leak Potential PN and LN | $E_L$ | -60mV |
| Leak Potential KC and APL | $E_L$ | -60mV |
| Threshold Potential ORN and KC | $V_T$ | -35mV |
| Threshold Potential PN and LN | $V_T$ | -30mV |
| Threshold Potential APL | $V_T$ | -30mV |
| Resting Potential ORN and LN | $V_r$ | -60mV |
| Resting Potential PN | $V_r$ | -60mV |
| Resting Potential KC | $V_r$ | -55mV |
| Resting Potential APL | $V_r$ | -60mV |
| Refractory Time | $\tau_{ref}$ | 2ms |
| Synaptic Parameters |  |  |
| Excitatory Potential | $E_E$ | 0mV |
| Inhibitory Potential | $E_I$ | -75mV |
| Excitatory Time Constant | $\tau_e$ | 5ms |
| Inhibitory Time Constant | $\tau_i$ | 10ms |
| Synaptic Weights |  |  |
| Weight Input-ORN | wORNinputORN | 3nS |
| Weight ORN-PN | wORNPN | 30nS |
| Weight ORN-LN | wORNLN | 9nS |
| Weight LN-PN | wLNPN | 2nS |
| Weight PN-KC | wPNKC | 1nS |
| Weight KC-APL | wKCAPL | 50nS |
| Weight APL-KC | wAPLKC | 100nS |

**Table 1. Network simulation parameters.**
|  |  |  |
| --- | --- | --- |
| Adaptation Time Constant | $\tau_{Ia}$ | 1000ms |
| Adaptation Reversal Potential | $E_{Ia}$ | -90mV |

All code for the software implementation is accessible via https://github.com/nawrotlab/DrosophilaOlfactorySparseCoding

### Spontaneous activity

The input to the ORNs in our network model was modeled as stochastic point process realizations. It mimicks the sum of spontaneous receptor activation and odor driven activation of the ORNs. On the chip, each ORN received a Poisson input with 5 Hz baseline intensity. In the simulation each ORN received excitatory synaptic input modelled as a gamma process (shape parameter k= 3) with 5 Hz baseline intensity. The spontaneous firing rate of larval ORNs was previously measured in the range of 0.2 – 7.9 Hz, depending strongly on receptor type and odor identity [67, 68]. On the chip we measured a spontaneous ORN firing rate of 6.2 ± 3.0Hz. In the simulated model the average ORN baseline activity was estimated as 6.0 ± 1.4 Hz. Thus, ORNs on chip and in the simulation exhibit a similar spontaneous activity in the upper range of the empirical distribution.

### Odor stimulation protocol

On the chip and in the computer simulation we included a warm up time (1.5 s and 0.3 s, respectively), which was excluded from the analyses. On the chip this restored the baseline biases following odor application. In the computer simulation this warm up period ensured that neuronal membranes and conductances were more heterogeneous at the beginning of the experiments.

We used a set of three different odors to study the effect of odor similarity (Fig. 1 B, upper panel). Fig 1B shows the activation profile (point process intensities) and overlap of all three odors across the 21 input channels. For each odor, the profile indicates the ORN-type specific activation level, mimicking the fact that each ORN expresses a genetically different receptor type. Similarity of odors is represented in the overlapping activation where odor 1 and odor 3 are distant (zero overlap), while odor 2 is constructed to have the same amount of overlap with the two other odors. The stimulation protocol assumes a 2 s odor stimulus on top of the 5 Hz baseline input with an activation rate according to Fig. 1 B.

### Data Analysis

#### Sparseness measure

Sparseness was quantified by the widely used modified version [69] of the Treves–Rolls measure [70]

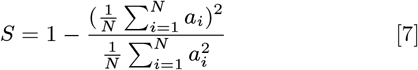

where *a_i_* indicates either the spike count of neuron *i* (population sparseness, *S_pop_),* or the binned (Δ*t* = 20 ms) population spike count (temporal sparseness, *S_tmp_*) for the 2 s with odor stimulation. *S* assumes values between zero and one, with high values indicating sparse responses. Both measures were averaged over 20 trials. To test the effect of excluding specific sparseness mechanisms from the model the combined data from all three odors was compared. To test for significance of the effects of lateral inhibition and SFA, the condition with only lateral inhibition enabled was compared with the condition with only SFA present (LN vs. SFA) using a t-test for related samples. To test the effect of feedback inhibition via the APL, the condition including all mechanisms (LN+APL+SFA) was compared with LN+SFA. Tests were performed independently for temporal and population sparseness.

#### Activation measure

We define an additional measure of activation as

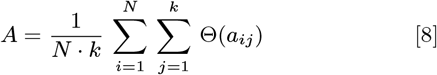

where *a_ij_* indicates the spike count of neuron *i* in the time bin *j* and Θ is the Heaviside step function. Thus, Θ(*a_ij_*) indicates the binary response of neuron *i* in time bin *j*. Tc asses population activation *A_pop_* we apply a single time bin foi the complete 2 s odor stimulation time. Then *A_pop_* measures the fraction across all *N* neurons that are odor-activated by a1 least one single spike. We quantify temporal activation *A_tmp_* by binning the stimulus time into *k* = 20 bins of *w =* 100 ms Thus, *A_tmp_* measures the binary response probability across time bins for each neuron. Both measures were then averaged over 20 trials and three odors. Our definition of activation is related to the complementary measure of ’activity sparseness’ defined in [69].

#### Distance measure

To assess the differences in odor distance between sparse and dense KC odor code we used the cosine distance (Eqn. 9). Vectors a and b each represent the average number of spikes evoked by all 72 KCs during the two second odor presentation across 20 independent model instances. Cosine distance between a and b was calculated as:

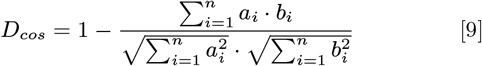

#### Correlation across sparseness conditions

To test for qualitatively similar effects of the different sparseness conditions on the chip and simulation we correlated the results across the six data points (sparseness conditions) between the chip and the simulation. For significance testing we generated 100 random unique permutations of the means from the simulation and correlated these 100 data series with that of the chip (LN+APL+SFA,LN+APL,APL+SFA,LN+SFA,LN,SFA; Fig. 3). The average of these 100 correlations was 0.07 (*sd* = 0.42) for *S_pop_* and 0.01 (*sd* = 0.52) for *S_tmp_* with *Q*_0.75_ = 0.37 and *Q*_0.75_ = 0.28. In both cases the distribution of correlations was normal, established using D’Agostino-Pearson test for normality. The average of these 100 correlations each was used to evaluate the similarity of the effects on the chip and in the simulation.

## Results

The larval nervous system with its limited neural network size and low complexity lends itself to the emulation on neuromorphic hardware. We analyzed a single hemisphere olfactory network model of the first instar *Drosophila* larva with <200 neurons and <1000 synapses comparing an implementation on the neuromorphic hardware DYNAP-SE [60] with a computer simulation of the same network. We were particularly interested in the contribution of different cellular and circuit mechanisms to the transformation of a dense input pattern at the periphery into a sparse odor representation in the MB.

### Olfactory pathway model

Our spiking neural network model comprises four computational layers (Fig. 1 A). Its structure, the size of the neuron populations and their connectivity are based on the exact connectome of a single hemisphere as reconstructed from electron-microscopic data of one individual *Drosophila larva* by Eichler and colleagues [62]. Peripheral processing is carried out by 21 ORNs, each expressing a different olfactory receptor type [71]. ORNs make one-to-one excitatory connections with 21 PNs and with 21 LNs that together constitute the antennal lobe. Each LN forms inhibitory synapses onto all PNs, establishing lateral inhibition. The PNs make divergent random connections with a total of 72 KCs, the primary cells of the mushroom body, where each KC receives excitatory input from 1-6 PNs. The APL receives input from all of the matured KCs [63]. All KCs with a well-developed dendrite [62] fall into this category and those are the only ones included in our circuit model. We therefore assume a dense convergent connectivity with essentially all presynaptic KCs (in our case 64 out of 72 due to technical limitations on the chip, see Methods). We further implemented inhibitory feedback from the APL onto all KCs [63]. Overall, this blueprint of the olfactory network is highly similar to that in the adult fly except for the smaller neuron numbers and reduced anatomical complexity (see Discussion). Each model instance implemented here utilizes the exact same connectivities. We thus simulate a single individual rather than an average animal.

### Circuit motifs and cellular adaptation

Our network model utilizes different cellular and circuit mechanisms that have been suggested to support a sparse code in the insect mushroom body. To this end, the network topology includes three relevant motifs. First, the LN connectivity in the antennal lobe constitutes lateral inhibition as a motif that generally enhances neural contrast [34, 72] and that is implemented in the olfactory system of virtually all insects [36, 73–78], as well as in computational models thereof [25, 28, 52, 79]. Second, the random connectivity from PNs to a larger number of KCs is net divergent and sparse, expanding the dimensionality of the coding space [51, 80, 81]. Third, our model includes inhibitory feedback from the APL neuron onto all KCs. This has been shown to directly affect KC populations sparseness in the adult fly [47, 82] (see Discussion).

At the cellular level, all neurons in the network are modeled as leaky integrate-and-fire neurons. ORNs and KCs are equipped with a cellular SFA mechanism, a fundamental and ubiquitous mechanism in spiking neurons [34, 83]. ORNs have been shown to adapt during ongoing stimulation *in vivo*, both in larval [84] and adult [85, 86] *Drosophila.* The exact nature of the adaptation mechanism in the ORNs is still under investigation [85, 87, 88]. In KCs, a strong SFA conductance has conclusively been demonstrated in the cockroach [89] and the bee [90].

### Dynamics of network response to odor stimulation

The response dynamics across all network stages to a single constant odor stimulation (Fig. 1 B) with odor 1 is shown in Fig. 2 A (chip) and Fig. 2B (simulation). At stimulus onset, a subset of all ORNs is activated according to the corresponding receptor response profile (Fig. 1B, top). The ORN responses are phasic-tonic as a result of SFA. The spike count histogram averaged across the 21 neurons of the ORN population fits the typical experimentally observed response profile observed in adult *Drosophila* [85, 91]. In the larva, little is known about stimulus adaptation in the ORNs [68]. The physical realization of SFA on chip is different from the simulation, which may partly explain the delayed response to odor onset- and offset of some neurons and the initially slower increase of the phasic response on the chip (Fig. 2 A, see Discussion). The off-response expressed in a prolonged silence of the odor-activated ORNs in the simulation is an effect of SFA: The integrated adaptation current that has reached a steady state during the odor stimulation period now decays only slowly, acting in a hyperpolarizing fashion and thus reducing spiking probability [52] of the ORNs. This effect is barely visible and delayed on the chip (see Discussion). At the level of the antennal lobe both PNs (dark blue) and LNs (light blue) are excited only by the ORNs and thus follow their phasic-tonic response behavior and exhibit an inhibited off-response (Fig. 2), although neither neuron type is adaptive itself. The spatio-temporal response pattern of the PNs and LNs resembles the typical response pattern measured *in vivo* in adult flies and bees [74, 92, 93], including a inhibitory off-response in many neurons [85, 94, 95].

**Fig. 2.**
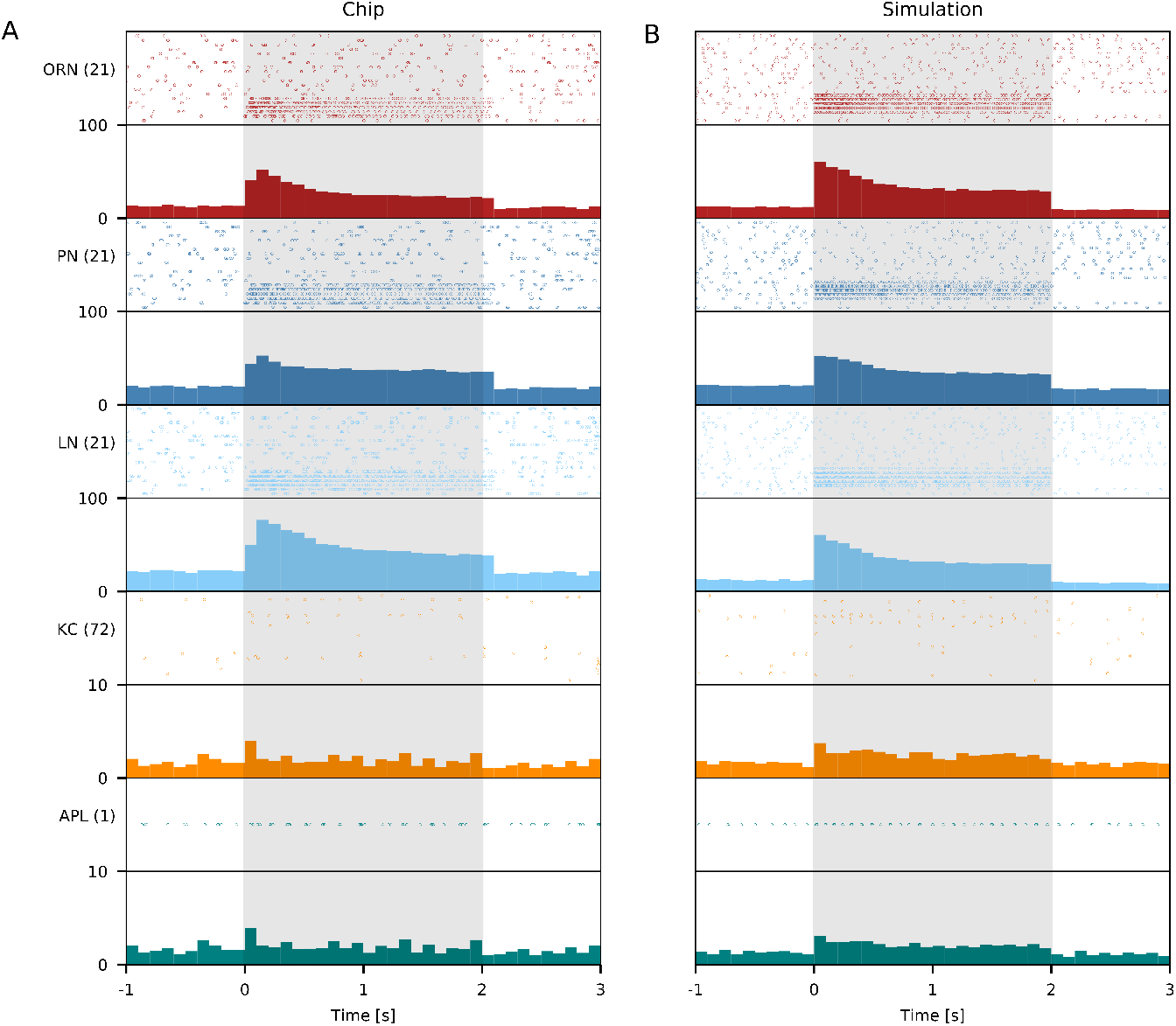
Dynamic network response. Network response to a stimulation with odor 1 for the chip (A) and the simulation (B). The odor was presented for 2s, preceded by a 2s baseline and followed by 2s again without odor. Warmup times are excluded and only the time window between 1 and 5 seconds is shown here. Odor onset is at time=0 and odor presence in the stimulation protocol is indicated by the shaded area throughout. Each dot denotes a single spike event of the respective neuron during an individual exemplary experiment. The lower panel (A,B) for each neuron population displays the averaged population spike count (across 20 trials) with a bin width of 100ms.

The KCs show very little spiking during spontaneous activity on the chip and in simulation. Only very few KCs do respond to odor stimulation (population sparse response) with only a single or few spikes (temporally sparse response). Spontaneous activity and response properties match well the *in vivo* situation as observed in various species [36, 54, 58]. The population spike count indicates a very brief population response within the first 100 ms, while the tonic KC response remains only slightly above the spontaneous activity level (cf. [54]. Finally, the single APL driven by the excitatory KC population follows the brief phasic and weak tonic response of the KCs.

### Analysis of sparsening factors in space and time

We investigate the translation from the peripheral dense code in the ORN and PN population into a central sparse code in the KC population, disentangling the contribution of the three fundamental biological mechanisms: cellular adaptation (SFA), lateral inhibition in the AL, and feedback inhibition in the MB. We systematically varied the composition of the three mechanisms in our network, yielding five different conditions (Fig. 3) in which either one or two mechanisms were deactivated. SFA was only deactivated at the KC level and still present in ORNs. We did not vary the PN-KC connectivity pattern as this is identical to the anatomical pattern reported for the individual animal that we used as a reference.

**Fig. 3.**
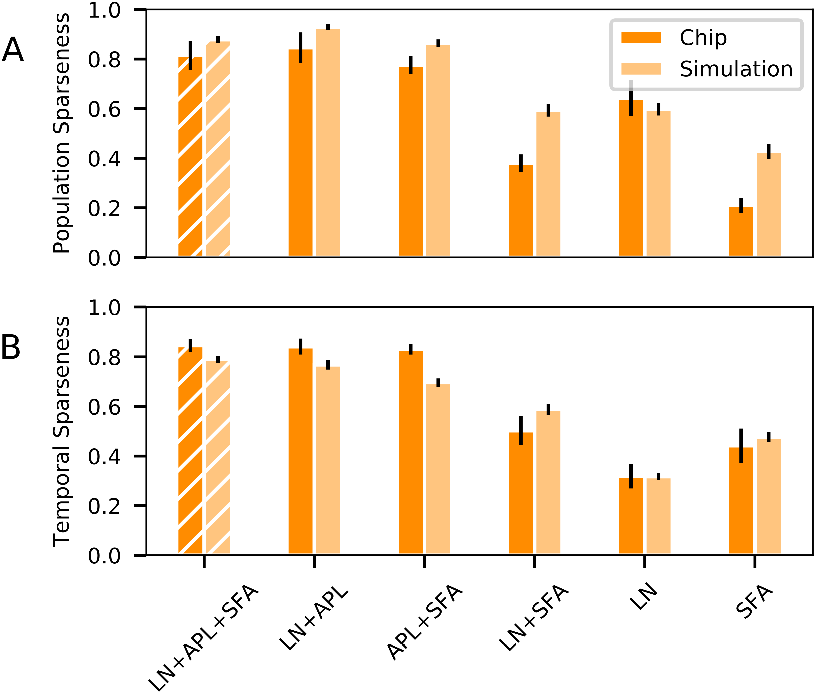
Mechanisms underlying population and temporal sparseness. (A) Comparison of KC population sparseness *S_pop_* between the chip and the simulation during 2s with odor stimulation. The data is averaged over 20 experiments and three odors (error bars denote standard deviations). Six experimental conditions were tested with a different set of sparseness mechanisms enabled. The respective mechanisms are listed below with LN (lateral inhibition via local interneurons), SFA (spike frequency adaptation) and APL (feedback inhibition via the APL). E.g. ‘LN+SFA’ denotes the presence of SFA and lateral inhibition. (B) Temporal sparseness *S_tmp_*was computed for the same set of conditions.

We additionally quantified the population activation by measuring the fraction of stimulus-activated KCs across the different conditions (see Methods) and find that it depends on the the sparseness mechanisms. It is lowest in the control condition with 20.6 (28.6 %) responding neurons on the chip and 16.7 (22.9%) in the simulation (see Supplements, Fig. S1 A). Our results show that APL is the single crucial mechanism necessary for establishing a high population sparseness in our model. All conditions that lack feedback inhibition show strongly reduced values of *S_pop_*. Lateral inhibition can to some degree recover sparseness on the chip and in the simulation.

We now consider temporal sparseness, which again reached high values in the control condition on the order of *S_tmp_* ≈ 0.8 (hatched bars in Fig. 3B). Comparing the different conditions we find that APL feedback inhibtion and SFA in the KCs have a strong supporting effect for temporal sparseness. Any condition that involves the APL reached similar high values for *Stmp*. Without the APL, SFA can partially ensure temporal sparseness on the chip and in the simulation. This is also reflected in the temporal activation measure that computes the fraction of active time bins (of 100 ms duration) for the complete 2s stimulation time (see Methods). The results shown in Fig. S1 B mirror our results in Fig. 3 B. In the sparse control condition KCs are active in on average only 2.3 % and 3.4 % of the response bins for the chip and the simulation, respectively.

Overall, we observed the same mechanistic effects on chip and in the simulation for the different combinations of activated and inactivated mechanisms (Fig. 3). The pattern of sparseness values across all six conditions is highly and significantly correlated between the chip and the simulation results, both for *S_pop_* (*r* = 0.94) and *S_tmp_* (*r* = 0.96) in comparison with the correlation of randomly permuted pattern of sparseness values (see Methods) with maximum correlations of 0.91 and 0.87 for *S_pop_* and *S_tmp_* across 100 permutations, respectively.

### Sparse representation supports stimulus separation

How does the encoding of different odors at the KC level compare between the sparse control condition and a non-sparse condition? Feedback and lateral inhibition supported population sparseness in the KC population. We thus compared the control condition to the network in which both inhibitory mechanisms were disabled by quantifying the pairwise distance between KC stimulus response patterns for any two different odors. Fig. 4 shows the response rates averaged over the 2 s stimulus duration for the three different stimuli for both chip Fig. 4 A,B,C and simulation Fig. 4 C,D,E. Only a fraction of the KCs responded to any odor (*S_pop_* > 0.8, Fig. 3 B,E) in the sparse condition. However, when feedback and lateral inhibition are disabled, essentially all KCs showed an odor response to any of the three odors (Fig. 4 C,F).

**Fig. 4.**
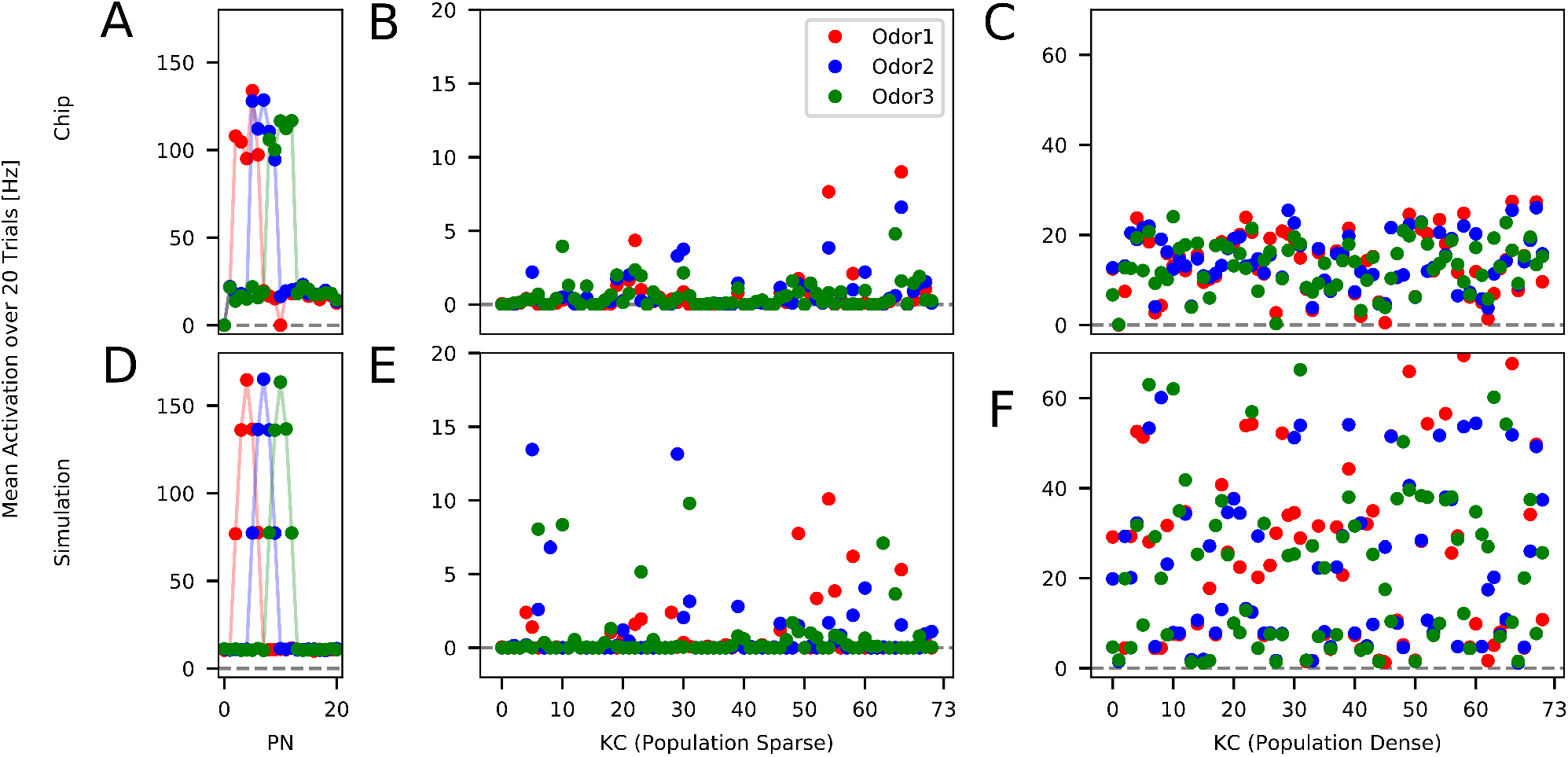
Response pattern overlap. Average spike frequency (over 20 trials) for Chip (A,B,C) and simulation (D,E,F) in response to three different odors. The odors were presented for 2s (for information on the experimental protocol please refer to Figure. 1 B). All panels display the overlap between the different odor representations either on PN level (A,D), KC (B,E) and non-population-sparse KC in a condition with only SFA enabled (C,F). Overlap indicates a low ability to differentiate between odors.

A similar result is obtained when looking at cosine distances between KC odor representations. Independently of the odor identities, average pairwise cosine distance was considerably larger in the sparse condition (chip: 0.39(0.2); simulation: 0.85(0.06)) than in the non-sparse (SFA only) condition (chip: 0.07(0.02); simulation: 0.31(0.09)), indicating a similar effect of population sparseness on odor discriminability on the chip and in the simulation.

## Discussion

In the present manuscript we addressed two major questions. First, we asked whether the re-coding from a dense peripheral olfactory code into a sparse central brain representation of odors can be achieved in the small spiking neural network model of *Drosophila* larva. To this end we tested the relevance of three fundamental mechanisms:

- cellular adaptation
- lateral inhibition
- feedback inhibition

in establishing population and temporal sparseness. Second, we explored the feasibility of applying this coding scheme on real-time analogue neuromorphic hardware by comparing hardware implementation with software simulation at the relevant levels of stimulus encoding and processing.

### Neuromorphic implementation versus computer simulation

While the software uses identical parameters for all neurons and synapses for any given population, there is considerable heterogeneity across the physical hardware implementation, in particular of conductances and capacities [4, 96, 97]). This heterogeneity manifests e.g. in spiking thresholds, postsynaptic current amplitudes and time constants of the neuron membrane, postsynaptic currents, and SFA currents. This generally matches the biological heterogeneity.

Our results show that the on-chip implementation achieved the transformation from dense to sparse coding in space and time despite the parameter heterogeneity in the small neuron populations. We obtained the same general results for our network analyses on hardware and in simulation albeit with small differences. These may arise from the fact that there is no one-to-one correspondence of the biophysical neuron parameters in the software simulation (Table 1) and the set points of the hardware electronic circuits. While parameter setting is straight forward in the computer simulation, it requires the adjustment of biasing currents on the chip. A particular challenge was the adjustment of the SFA time constant on the chip that required the post hoc estimate of the effective adaptation time constant. In our simulations a difference in the SFA effect becomes evident in the delayed SFA on- and offset effects of the chip.

There are a number of advantages and disadvantages in using the specific hardware solution tested here. The fact that the DYNAP-SE [60] operates in real-time makes it suitable for the spiking control of autonomous robots [98, 99] and renders computational speed independent of network size. Even for the small larval network considered here (exactly 136 neurons and 833 synapses) simulations were several times slower than real-time with 3.8 s simulation time per 1 s biological time at a resolution of 0.1 ms (single core CPU, 64 bit PC, Ubuntu 18.04.5). Simulation time can be sped up to meet real-time demand even for large network sizes on specialized systems [100, 101].

A challenge with the DYNAP-SE [60] board was the (thermal) instability of the bias currents used to adjust parameters. This lead to parameter drifts over the course of hours, which made it necessary to complete a set of experiments within a limited time range to ensure comparability of the results. Before the next set of experiments, biases had to be re-adjusted (in particular for the adaptation circuit). Thus, each set of experiments implied an individual model instance that may be compared to an individual animal. Still, the variability across model instances was only slightly larger on the chip than in the simulation (Fig. 3). In addition, new neuromorphic circuit designs will be able to compensate for these drifts by using proper *temperature compensated* bias generator circuits [102].

### Mechanisms and function of population sparseness

Population sparseness at the KC level has been demonstrated for a number of species in the adult stage (see Introduction). Our model based on the current knowledge of anatomical structures within the *Drosophila* larva olfactory pathway suggests that population sparseness is already implemented in the larva with similar functional benefits. Different mechanisms have been suggested for the generation of population sparseness. A fundamental anatomical basis for a sparse code is the sparse and divergent connectivity between PNs and a much larger population of KCs [37, 52]. Each KC receives input from only a few PNs and thus establishes a projection from a lower into a higher dimensional space, ideally suited to generate distinct activity patterns that foster associative memory formation. Additionally, there is evidence for a low excitability of the individual KCs that require collective input from several PNs to be activated [36, 37, 89]. Connectivity in our model is based on the exact numbers from electron microscopic reconstructions of neurons and synapses in one individual brain [62]. We did not attempt to adjust excitability of KCs or PN::KC connection strength for optimal population sparseness.

Feedback inhibition has repeatedly been suggested to underlie population sparseness in several animals, including the fly larva [103]. Empirical evidence has been provided in particular in bees [94, 104] and adult flies [47, 82]. Several modeling studies have used feedback inhibition to support a sparse KC population code in larger adult KC populations [25, 27, 28, 105]. Indeed, our study shows that inhibitory feedback from the single APL neuron effectively implements a sparse code in the small population of 72 KCs (Fig. 3 A). We chose to model the APL as a spiking neuron that receives input solely from KCs and inhibits KCs in a closed loop. This decision was based on experimental evidence indicating a clear polarity of the APL with input in the MB lobes and pre-synaptic densities in the calyx, presumably onto the KC dendrites [63]. Whether the APL neuron generates sodium action potentials, however, is not clear in the larva [103] and has been challenged in the adult [47]. In addition, inhibitory feedback connections within the mushroom body have been implicated in learning through inhibitory plasticity in bees and flies, thereby modulating the sparse KC population code [47, 104, 105].

As a third factor, lateral inhibition within the *Drosophila* antennal lobe has been shown to increase population sparseness at the KC level [91, 106] and in a model thereof [52]. This model study showed a strong effect of lateral inhibition on population sparseness in a network tuned to the anatomy of the adult fly. In the present larval model we found a supportive effect. With lateral inhibition alone the model reached *S_pop_* ≈ 0.6. The interplay of feedback inhibition and lateral inhibition boosted population sparseness to *S_pop_* ≈ 0.8 (Fig. 3 A). This observation is different from our previous results in a network simulation modelled after the adult fly [28] where lateral inhibition in the AL was sufficient to implement a high population sparseness and APL feedback inhibition had a mainly supporting effect. The fact that lateral inhibition is less effective in the larval than in the adult Drosophila model [52] is thus likely due to the one-to-one connectivity between the 21 ORNs and 21 PNs in the larva, which requires very strong excitatory synapses. This specific configuration establishes a dominant feed-forward component in the larval olfactory pathway (Fig. 1 A).

Sparse stimulus representation across the neuronal population supports minimal overlap of and correlation across stimulus-specific spatial response patterns [34, 52, 82, 107, 108], which in turn benefits associative memory formation and increases memory capacity [47, 53, 70, 109]. We confirmed an increased inter-stimulus distance in the KC coding space on the chip and in the simulation when all sparseness mechanisms take effect.

### Mechanisms and function of temporal sparseness

Temporal sparseness in the insect MB has been physiologically described in various species. It is expressed in a highly phasic stimulus response that typically consists of only a single or very few spikes and that is temporally locked to stimulus onset or to a fast transient increase in stimulus amplitude while the tonic stimulus response is almost absent [36, 54, 58]. In our model we implemented two mechanisms that can support temporal sparseness, inhibitory feedback via the spiking APL neuron and SFA. Our analysis in Fig. 3 B showed that inhibitory feedback has the strongest effect, confirming experimental [110, 111] and modeling results [25, 27, 28, 112]. Cellular adaptation (SFA) showed a smaller but supporting effect in our network, which is partially in line with our previous models of the adult fly [28, 52, 110] in which we showed that SFA alone can suffice to generate high temporal sparseness.

Importantly, cellular adaptation has additional effects on stimulus coding that are not analyzed here. Being a self-inhibiting mechanism it reduces overall spiking activity, contributing to the low spontaneous and response rates in the KC population that has been repeatedly documented in various insect species [36, 54, 58]. Moreover, SFA leads to a regularization of the neuron’s spike output and a reduction of the trial-to-trial variability, effectively improving response reliability [110, 113]. Finally, SFA introduces a short-term stimulus memory expressed in the conductance state of the excited neuron population, which decays with the SFA time constant [52].

Temporal sparseness was influenced strongly by SFA in the KCs and by recurrent feedback inhibition. It usually shows as longer inter-spike-intervals both in physiological data [36, 54, 94] as well as in modeling results [28, 52, 110]. Besides the prolongation of the inter-spike-intervals over the entire duration of the experiment, SFA also caused the commonly observed odor onset effect [36, 54, 58, 94, 111] in ORNs and KCs. In our data this effect was somewhat concealed in the KCs by the overall small number of spike responses. This is a tribute to the biological plausibility with respect to data collected from adult *Drosophila*, where the KC rarely show spikes at baseline [58] and a very sparse odor response pattern [58, 114]. Due to the SFA in the ORN population that was active in all experimental conditions there was a good degree of temporal sparseness in the LN-only condition as well (especially on the chip). Again we chose to accept this effect as a baseline level of sparseness to compare other conditions against. In both implementations the expected effects of SFA in the KCs could be observed.

We have previously argued that the major functional role of temporal sparseness is the rapid and reliable stimulus encoding in a temporally dynamic environment [28, 110]. Indeed, temporal dynamics is high in the natural olfactory environment and depends on air movement and on animal speed, the latter being particularly high in flying insects. As a result, adult insects during flight or locomotion may encounter a rapid on-off stimulus scenario when passing through a thin odor filament [115–119]. It remains an open question whether the SFA mechanism is at all present in the KCs during larval stages and electrophysiological approaches to neural coding in the larva is scarce. Representation of high temporal stimulus dynamics is likely of minor importance for the larva as its locomotion is slow and the natural environment suitable for larval development such as e.g. a rotten fruit likely provides little olfactory dynamics. However, larva do perform chemotaxis and thus are able to sample olfactory gradients.

### Outlook

Our current research extends the present model towards a plastic spiking network model of the larva that can perform associative learning and reward prediction [120] inspired by recent modelling approaches in the adult [121, 122]. Together with biologically realistic modeling of individual larva locomotion and chemotactic behavior [16] this will allow to reproduce behavioral [123–127] and optophysiological observations [63, 128, 129] and to generate testable hypothesis at the physiological and behavioral level. In the future this may inspire modeling virtual larvae exploring and adapting to their virtual environment in a closed loop scenario and the implementation of such mini brains on compact and low-power neuromorphic hardware for the spiking control of autonomous robots [7, 28, 130, 131].

## ACKNOWLEDGMENTS

This projects is funded by the German Research Foundation (DFG) within the Research Unit ‘Structure, Plasticity and Behavioral Function of the *Drosophila* mushroom body’(DFG-FOR 2705, grant no. 403329959, https://www.uni-goettingen.de/en/601524.html), and in part by the European Research Council (ERC) under the European Union’s Horizon 2020 research and innovation program grant agreement No 724295 (NeuroAgents). AMJ received additional support from the Research Training Group ‘Neural Circuit Analysis’ (DFG-RTG 1960, grant no. 233886668). The authors would like to acknowledge the financial support of the CogniGron research center and the Ubbo Emmius Funds (Univ. of Groningen). We thank Sören Rüttger for support with the neuromorphic hardware setup, and Hannes Rapp, Panagiotis Sakagiannis and Bertram Gerber for valuable discussions.

## Supplements

**Fig. S1.**
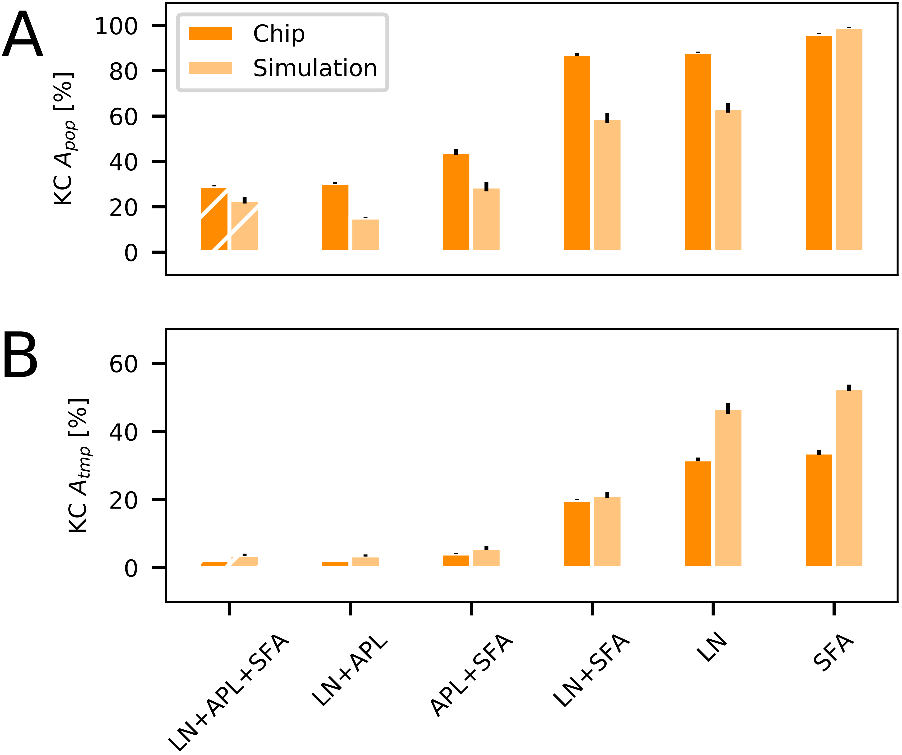
Population activation and temporal activation. (A) Comparison of KC population activation *A_pop_* between the chip and the simulation during the 2 s odor stimulation, averaged over 20 trials and three odors (error bars denote standard deviation across odors). Six experimental conditions were tested, each with a different set of sparseness mechanisms enabled. The respective mechanisms are listed below with LN (lateral inhibition via local interneurons), SFA (spike frequency adaptation) and APL (feedback inhibition via the APL). E.g. ‘LN+SFA’ denotes the presence of SFA and lateral inhibition. (B) Temporal activation *A_tmp_* was computed for the same set of conditions by averaging the number of 100 ms time windows during which each KC is active. The binary activation measure is complementary to the sparseness measure in main Fig. 3.

**Fig. S2.**
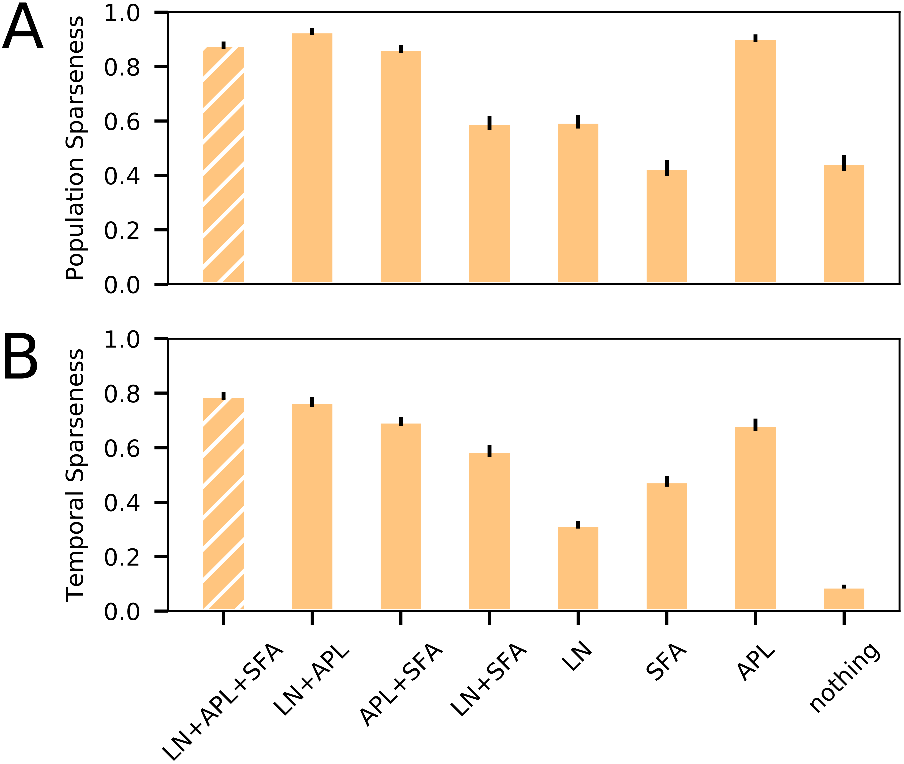
Mechanisms underlying population and temporal sparseness. (A) KC population sparseness of the simulation during 2s with odor stimulation. The data is averaged over 20 experiments and three odors (error bars denote standard deviations across odors). In the simulation two more experimental conditions were tested in addition to the those shown in main Fig. 3. The respective mechanisms are listed below with LN (lateral inhibition via local interneurons), SFA (spike frequency adaptation) and APL (feedback inhibition via the APL). E.g. ‘LN+SFA’ denotes the presence of SFA and lateral inhibition. (B) Temporal sparseness was computed for the same set of conditions.

## Notes

### Competing Interest Statement

The authors have declared no competing interest.

## References

1. C Mead, Neuromorphic Electronic Systems. Proc. IEEE 78, 1629–1636 (1990).

2. G Indiveri, Y Sandamirskaya, The importance of space and time for signal processing in neuromorphic agents. IEEE Signal Process. Mag. 36, 16–28 (2019).

3. E Neftci, et al., Synthesizing cognition in neuromorphic electronic systems. Proc. National Acad. Sci. United States Am. 110, 3468–3476 (2013).

4. M Schmuker, T Pfeil, MP Nawrot, A neuromorphic network for generic multivariate data classification. Proc. National Acad. Sci. United States Am. 111, 2081–2086 (2014).

5. A Diamond, T Nowotny, M Schmuker, Comparing Neuromorphic Solutions in Action: Implementing a Bio-Inspired Solution to a Benchmark Classification Task on Three Parallel-Computing Platforms. Front. Neurosci. 9, 491 (2016).

6. B Cramer, et al., Training spiking multi-layer networks with surrogate gradients on an analog neuromorphic substrate. https://arxiv.org/abs/2006.07239 (2020).

7. LI Helgadottir, J Haenicke, T Landgraf, R Rojas, MP Nawrot, Conditioned behavior in a robot controlled by a spiking neural network in International IEEE/EMBS Conference on Neural Engineering, NER. pp. 891–894 (2013).

8. F Galluppi, et al., Event-based neural computing on an autonomous mobile platform in 2014 IEEE International Conference on Robotics and Automation (ICRA). (IEEE), pp. 2862–2867 (2014).

9. T Schoepe, et al., Finding the gap: Neuromorphic motion vision in cluttered environments. https://arxiv.org/pdf/2102.08417.pdf (2021).

10. M Heisenberg, Pattern recognition in insects. Curr. Opin. Neurobiol. 5, 475–481 (1995).

11. D Carrasco, MC Larsson, P Anderson, Insect host plant selection in complex environments. Curr. Opin. Insect Sci. 8, 1–7 (2015).

12. M Laska, CG Galizia, M Giurfa, R Menzel, Olfactory discrimination ability and odor structureactivity relationships in honeybees. Chem. Senses 24, 429–438 (1999).

13. G Meckenhäuser, S Krämer, F Farkhooi, B Ronacher, MP Nawrot, Neural representation of calling songs and their behavioral relevance in the grasshopper auditory system. Front. Syst. Neurosci. 8, 183 (2014).

14. A Avarguès-Weber, G Portelli, J Benard, A Dyer, M Giurfa, Configural processing enables discrimination and categorization of face-like stimuli in honeybees. J. Exp. Biol. 213, 593–601 (2010).

15. M Collett, L Chittka, TS Collett, Spatial memory in insect navigation. Curr. Biol. 23, 789–800 (2013).

16. A Wystrach, P Graham, What can we learn from studies of insect navigation? Animal Behav. 84, 13–20 (2012).

17. R Menzel, U Greggers, The memory structure of navigation in honeybees. J. Comp. Physiol. A 201, 547–561 (2015).

18. M Knaden, P Graham, The sensory ecology of ant navigation: from natural environments to neural mechanisms. Annu. review entomology 61, 63–76 (2016).

19. P Sakagiannis, AM Jürgensen, MP Nawrot, A realistic locomotory model of drosophila larva for behavioral simulations. https://www.biorxiv.org/content/10.1101/2021.07.07.451470v1.abstract (2021).

20. L Chittka, M Giurfa, JA Riffell, The mechanisms of insect cognition. Front. psychology 10, 2751 (2019).

21. M Dacke, MV Srinivasan, Evidence for counting in insects. Animal cognition 11, 683–689 (2008).

22. P Skorupski, H MaBouDi, HS Galpayage Dona, L Chittka, Counting insects. Philos. Trans. Royal Soc. B: Biol. Sci. 373, 20160513 (2018).

23. SR Howard, A Avarguès-Weber, JE Garcia, AD Greentree, AG Dyer, Numerical ordering of zero in honey bees. Science 360, 1124–1126 (2018).

24. H MaBouDi, et al., Bumblebees use sequential scanning of countable items in visual patterns to solve numerosity tasks. Integr. comparative biology 60, 929–942 (2020).

25. C Assisi, M Stopfer, M Bazhenov, Optimality of sparse olfactory representations is not affected by network plasticity. PLoS computational biology 16, e1007461 (2020).

26. J Wessnitzer, JM Young, JD Armstrong, B Webb, A model of non-elemental olfactory learning in Drosophila. J. computational neuroscience 32, 197–212 (2012).

27. P Ardin, F Peng, M Mangan, K Lagogiannis, B Webb, Using an insect mushroom body circuit to encode route memory in complex natural environments. PLoS computational biology 12, e1004683 (2016).

28. H Rapp, MP Nawrot, A spiking neural program for sensorimotor control during foraging in flying insects. Proc. National Acad. Sci. 117, 28412–28421 (2020).

29. J Müller, M Nawrot, R Menzel, T Landgraf, A neural network model for familiarity and context learning during honeybee foraging flights. Biol. Cybern. 112, 113–126 (2018).

30. T Rost, H Ramachandran, MP Nawrot, E Chicca, A neuromorphic approach to auditory pattern recognition in cricket phonotaxis in 2013 European Conference on Circuit Theory and Design (ECCTD). (IEEE), pp. 1–4 (2013).

31. T Dalgaty, E Vianello, B De Salvo, J Casas, Insect-inspired neuromorphic computing. Curr. opinion insect science 30, 59–66 (2018).

32. IA Lungu, A Riehle, MP Nawrot, M Schmuker, Predicting voluntary movements from motor cortical activity with neuromorphic hardware. IBMJ. Res. Dev. 61, 5–1 (2017).

33. T Dalgaty, JP Miller, E Vianello, J Casas, Bio-inspired architectures substantially reduce the memory requirements of neural network models. Front. neuroscience 15, 156 (2021).

34. HB Barlow, Sensory mechanisms, the reduction of redundancy, and intelligence in Mechanisation of thought processes. (Her Majesty’s Stationery Office, London), pp. 535–539 (1959).

35. BA Olshausen, DJ Field, Sparse coding of sensory inputs. Curr. Opin. Neurobiol. 14, 481–487 (2004).

36. J Perez-Orive, et al., Oscillations and sparsening of odor representations in the mushroom body. Science 297, 359–365 (2002).

37. RA Jortner, SS Farivar, G Laurent, A simple connectivity scheme for sparse coding in an olfactory system. The J. neuroscience: official journal Soc. for Neurosci. 27, 1659–1669 (2007).

38. LA Finelli, S Haney, M Bazhenov, M Stopfer, TJ Sejnowski, Synaptic Learning Rules and Sparse Coding in a Model Sensory System. PLOS Comput. Biol. 4, e1000062 (2008).

39. P Kloppenburg, MP Nawrot, Neural coding: Sparse but on time. Curr. biology 24, R957–R959 (2014).

40. Stopfer. Mark, Central processing in the mushroom bodies. Curr. Opin. Insect science 6, 99–103 (2015).

41. C Poo, JS Isaacson, Odor Representations in Olfactory Cortex: “Sparse” Coding, Global Inhibition, and Oscillations. Neuron 62, 850–861 (2009).

42. C Häusler, A Susemihl, MP Nawrot, Natural image sequences constrain dynamic receptive fields and imply a sparse code. Brain research 1536, 53–67 (2013).

43. J Wolfe, AR Houweling, M Brecht, Sparse and powerful cortical spikes. Curr. opinion neurobiology 20, 306–312 (2010).

44. JS Isaacson, Odor representations in mammalian cortical circuits. Curr. opinion neurobiology 20, 328–331 (2010).

45. T Hromádka, MR DeWeese, AM Zador, Sparse Representation of Sounds in the Unanesthetized Auditory Cortex. PLoS Biol. 6, e16 (2008).

46. SB Laughlin, TJ Sejnowski, Communication in neuronal networks. Science 301, 1870–1874 (2003).

47. AC Lin, AM Bygrave, A de Calignon, T Lee, G Miesenböck, Sparse, decorrelated odor coding in the mushroom body enhances learned odor discrimination. Nat. Neurosci. 17, 559–568 (2014).

48. A Litwin-kumar, et al., Optimal Degrees of Synaptic Connectivity Article Optimal Degrees of Synaptic Connectivity. Neuron 93, 1153–1164.e7 (2017).

49. SJ Caron, V Ruta, LF Abbott, R Axel, Random convergence of olfactory inputs in the Drosophila mushroom body. Nature 497, 113–117 (2013).

50. R Huerta, T Nowotny, M García-Sanchez, HD Abarbanel, MI Rabinovich, Learning classification in the olfactory system of insects. Neural computation 16, 1601–1640 (2004).

51. R Huerta, T Nowotny, Fast and robust learning by reinforcement signals: Explorations in the insect brain. Neural computation 21, 2123–2151 (2009).

52. R Betkiewicz, B Lindner, MP Nawrot, Circuit and cellular mechanisms facilitate the transformation from dense to sparse coding in the insect olfactory system. Eneuro 7, O.0305–18 (2020).

53. G Palm, Neural associative memories and sparse coding. Neural Networks 37, 165–171 (2013).

54. I Ito, RCY Ong, B Raman, M Stopfer, Sparse odor representation and olfactory learning. Nat. neuroscience 11, 1177–1184 (2008).

55. R Herikstad, J Baker, JP Lachaux, CM Gray, SC Yen, Natural movies evoke spike trains with low spike time variability in cat primary visual cortex. J. Neurosci. 31, 15844–15860 (2011).

56. B Haider, et al., Synaptic and Network Mechanisms of Sparse and Reliable Visual Cortical Activity during Nonclassical Receptive Field Stimulation. Neuron 65, 107–121 (2010).

57. G Laurent, M Naraghi, Odorant-induced oscillations in the mushroom bodies of the locust. J. Neurosci. 14, 2993–3004 (1994).

58. GC Turner, M Bazhenov, G Laurent, Olfactory representations by Drosophila mushroom body neurons. J. Neurophysiol. 99, 734–746 (2008).

59. R Menzel, The honeybee as a model for understanding the basis of cognition. Nat. Rev. Neurosci. 13, 758–768 (2012).

60. S Moradi, N Qiao, F Stefanini, G Indiveri, A Scalable Multicore Architecture with Heterogeneous Memory Structures for Dynamic Neuromorphic Asynchronous Processors (DYNAPs). IEEE Trans. on Biomed. Circuits Syst. 12, 106–122 (2018).

61. M Stimberg, R Brette, DF Goodman, Brian 2, an intuitive and efficient neural simulator. Elife 8, e47314 (2019).

62. K Eichler, et al., The complete connectome of a learning and memory centre in an insect brain. Nature 548, 175–182 (2017).

63. T Saumweber, et al., Functional architecture of reward learning in mushroom body extrinsic neurons of larval Drosophila. Nat. communications 9, 1104 (2018).

64. E Chicca, F Stefanini, C Bartolozzi, G Indiveri, Neuromorphic electronic circuits for building autonomous cognitive systems. Proc. IEEE 102, 1367–1388 (2014).

65. K Boahen, Communicating neuronal ensembles between neuromorphic chips in Neuromorphic Systems Engineering, ed. T Lande. (Kluwer Academic, Norwell, MA), pp. 229–259 (1998).

66. S Deiss, R Douglas, A Whatley, A pulse-coded Communications infrastructure for neuromorphic systems in Pulsed Neural Networks, eds. W Maass, C Bishop. (MIT Press), pp. 157–78 (1998).

67. DJ Hoare, et al., Modeling peripheral olfactory coding in drosophila larvae. PLoS One 6, e22996 (2011).

68. SA Kreher, JY Kwon, JR Carlson, The molecular basis of odor coding in the Drosophila larva. Neuron 46, 445–456 (2005).

69. B Willmore, DJ Tolhurst, Characterizing the sparseness of neural codes. Network: Comput. Neural Syst. 12, 255–270 (2001).

70. A Treves, ET Rolls, What determines the capacity of autoassociative memories in the brain? Network: Comput. Neural Syst. 2, 371–397 (1991).

71. A Couto, M Alenius, BJ Dickson, Molecular, anatomical, and functional organization of the Drosophila olfactory system. Curr. biology: CB 15, 1535–1547 (2005).

72. DH Hubel, TN Wiesel, Receptive fields of single neurones in the cat’s striate cortex. The J. physiology 148, 574–591 (1959).

73. JP Martin, et al., The neurobiology of insect olfaction: sensory processing in a comparative context. Prog. neurobiology 95, 427–447 (2011).

74. S Krofczik, R Menzel, MP Nawrot, Rapid odor processing in the honeybee antennal lobe network. Front. computational neuroscience 2, 9 (2009).

75. N Deisig, M Giurfa, JC Sandoz, Antennal Lobe Processing Increases Separability of Odor Mixture Representations in the Honeybee. J. Neurophysiol. 103, 2185–2194 (2010).

76. D Fuscà, P Kloppenburg, Odor processing in the cockroach antennal lobe—the network components. Cell Tissue Res. 383, 1–15 (2021).

77. B Leitch, G Laurent, Gabaergic synapses in the antennal lobe and mushroom body of the locust olfactory system. J. comparative Neurol. 372, 487–514 (1996).

78. SR Olsen, RI Wilson, Lateral presynaptic inhibition mediates gain control in an olfactory circuit. Nature 452, 956–960 (2008).

79. C Linster, BH Smith, A computational model of the response of honey bee antennal lobe circuitry to odor mixtures: Overshadowing, blocking and unblocking can arise from lateral inhibition. Behav. Brain Res. 87, 1–14 (1997).

80. T Nowotny, R Huerta, HD Abarbanel, MI Rabinovich, Self-organization in the olfactory system: One shot odor recognition in insects. Biol. Cybern. 93, 436–446 (2005).

81. TS Mosqueiro, R Huerta, Computational models to understand decision making and pattern recognition in the insect brain. Curr. opinion insect science 6, 80–85 (2014).

82. Z Lei, K Chen, H Li, H Liu, A Guo, The GABA system regulates the sparse coding of odors in the mushroom bodies of Drosophila. Biochem. Biophys. Res. Commun. 436, 35–40 (2013).

83. J Benda, Neural adaptation. Curr. Biol. 31, 110–116 (2021).

84. G Si, et al., Structured Odorant Response Patterns across a Complete Olfactory Receptor Neuron Population. Neuron 101, 950–962.e7 (2019).

85. KI Nagel, RI Wilson, Biophysical mechanisms underlying olfactory receptor neuron dynamics. Nat. neuroscience 14, 208–216 (2011).

86. S Gorur-Shandilya, M Demir, J Long, DA Clark, T Emonet, Olfactory receptor neurons use gain control and complementary kinetics to encode intermittent odorant stimuli. Elife 6, e27670 (2017).

87. C Martelli, JR Carlson, T Emonet, Intensity invariant dynamics and odor-specific latencies in olfactory receptor neuron response. The J. neuroscience: official journal Soc. for Neurosci. 33, 6285–6297 (2013).

88. SC Brandão, M Silies, C Martelli, Adaptive temporal processing of odor stimuli. Cell Tissue Res. 383, 1–17 (2021).

89. H Demmer, P Kloppenburg, Intrinsic Membrane Properties and Inhibitory Synaptic Input of Kenyon Cells as Mechanisms for Sparse Coding? J. Neurophysiol. 102, 1538–1550 (2009).

90. J Kropf, W Rössler, In-situ recording of ionic currents in projection neurons and Kenyon cells in the olfactory pathway of the honeybee. PloS one 13, e0191425 (2018).

91. RI Wilson, GC Turner, G Laurent, Transformation of olfactory representations in the drosophila antennal lobe. Science 303, 366–370 (2004).

92. V Bhandawat, SR Olsen, NW Gouwens, ML Schlief, RI Wilson, Sensory processing in the Drosophila antennal lobe increases reliability and separability of ensemble odor representations. Nat. Neurosci. 10, 1474–1482 (2007).

93. A Meyer, CG Galizia, MP Nawrot, Local interneurons and projection neurons in the antennal lobe from a spiking point of view. J. Neurophysiol. 110, 2465–2474 (2013).

94. P Szyszka, M Ditzen, A Galkin, CG Galizia, R Menzel, Sparsening and temporal sharpening of olfactory representations in the honeybee mushroom bodies. J. Neurophysiol. 94, 3303–3313 (2005).

95. MP Nawrot, Dynamics of sensory processing in the dual olfactory pathway of the honeybee. Apidologie 43, 269–291 (2012).

96. G Indiveri, et al., Neuromorphic silicon neuron circuits. Front. neuroscience 5, 73 (2011).

97. T Pfeil, et al., Effect of heterogeneity on decorrelation mechanisms in spiking neural networks: a neuromorphic-hardware study. Phys. Rev. X 6, 021023 (2016).

98. D Liang, et al., Neural State Machines for Robust Learning and Control of Neuromorphic Agents. IEEE J. on Emerg. Sel. Top. Circuits Syst. 9, 679–689 (2019).

99. Z Cao, et al., Spiking neural network-based target tracking control for autonomous mobile robots. Neural Comput. Appl. 26, 1839–1847 (2015).

100. JC Knight, T Nowotny, Larger gpu-accelerated brain simulations with procedural connectivity. Nat. Comput. Sci. 1, 136–142 (2021).

101. M Stimberg, DF Goodman, T Nowotny, Brian2genn: accelerating spiking neural network simulations with graphics hardware. Sci. reports 10, 1–12 (2020).

102. T Delbruck, R Berner, P Lichtsteiner, C Dualibe, 32-bit configurable bias current generator with sub-off-current capability in InternationalSymposium on Circuits andSystems, (ISCAS), 2010. (IEEE, IEEE, Paris, France), pp. 1647–1650 (2010).

103. LM Masuda-Nakagawa, K Ito, T Awasaki, CJ O’Kane, A single GABAergic neuron mediates feedback of odor-evoked signals in the mushroom body of larval Drosophila. Front. Neural Circuits 8, 1–11 (2014).

104. J Haenicke, N Yamagata, H Zwaka, M Nawrot, R Menzel, Neural correlates of odor learning in the presynaptic microglomerular circuitry in the honeybee mushroom body Calyx. eNeuro 5 (2018).

105. J Haenicke, Ph.D. thesis (2015) https://refubium.fu-berlin.de/handle/fub188/8123.

106. SR Olsen, V Bhandawat, RI Wilson, Divisive normalization in olfactory population codes. Neuron 66, 287–299 (2010).

107. D Marr, WT Thach, A Theory of Cerebellar Cortex in From the Retina to the Neocortex. (Birkhäuser Boston), pp. 11–50 (1991).

108. S Albus, A theory of cerebellar function. Math. Biosci. 10, 25–61 (1971).

109. T Nowotny, MI Rabinovich, R Huerta, HD Abarbanel, Decoding temporal information through slow lateral excitation in the olfactory system of insects. J. Comput. Neurosci. 15, 271–281 (2003).

110. F Farkhooi, A Froese, E Muller, R Menzel, MP Nawrot, Cellular adaptation facilitates sparse and reliable coding in sensory pathways. PLoS computational biology 9, e1003251 (2013).

111. A Froese, P Szyszka, R Menzel, Effect of GABAergic inhibition on odorant concentration coding in mushroom body intrinsic neurons of the honeybee. J. Of Comp. Physiol. A 200, 183–195 (2014).

112. T Kee, P Sanda, N Gupta, M Stopfer, M Bazhenov, Feed-Forward versus Feedback Inhibition in a Basic Olfactory Circuit. PLOS Comput. Biol. 11, e1004531 (2015).

113. F Farkhooi, E Muller, MP Nawrot, Adaptation reduces variability of the neuronal population code. Phys. Rev. E 83, 050905 (2011).

114. KS Honegger, RAA Campbell, GC Turner, Cellular-Resolution Population Imaging Reveals Robust Sparse Coding in the Drosophila Mushroom Body. J. Neurosci. 31, 11772–11785 (2011).

115. M Pannunzi, T Nowotny, Odor stimuli: Not just chemical identity. Front. physiology 10, 1428 (2019).

116. A Celani, E Villermaux, M Vergassola, Odor landscapes in turbulent environments. Phys. Rev. X 4, 041015 (2014).

117. M Kree, JÔ Duplat, E Villermaux, The mixing of distant sources. Phys. Fluids 25, 091103 (2013).

118. M Demir, N Kadakia, HD Anderson, DA Clark, T Emonet, Walking drosophila navigate complex plumes using stochastic decisions biased by the timing of odor encounters. Elife 9, e57524 (2020).

119. NJ Vickers, TA Christensen, TC Baker, JG Hildebrand, Odour-plume dynamics influence the brain’s olfactory code. Nature 410, 466–470 (2001).

120. AM Jürgensen, A Khalili, MP Nawrot, Reinforcement-mediated plasticity in a spiking model of the drosophila larva olfactory system in BMC Neuroscience2019. Vol. 20(Suppl 1): P225, p. 56 (2019).

121. JE Bennett, A Philippides, T Nowotny, Learning with reinforcement prediction errors in a model of the drosophila mushroom body. Nat. communications 12, 1–14 (2021).

122. M Springer, MP Nawrot, A mechanistic model for reward prediction and extinction learning in the fruit fly. Eneuro 8 (2021).

123. M Schleyer, D Miura, T Tanimura, B Gerber, Learning the specific quality of taste reinforcement in larval drosophila. elife 4, e04711 (2015).

124. B Gerber, T Hendel, Outcome expectations drive learned behaviour in larval Drosophila. Proc. Royal Soc. B: Biol. Sci. 273, 2965–2968 (2006).

125. M Schleyer, et al., The impact of odor-reward memory on chemotaxis in larval Drosophila. Learn. & memory (ColdSpring Harb. N.Y.) 22, 267–77 (2015).

126. T Saumweber, J Husse, B Gerber, Innate attractiveness and associative learnability of odors can be dissociated in larval Drosophila. Chem. Senses 36, 223–235 (2011).

127. M Schleyer, et al., A behavior-based circuit model of how outcome expectations organize learned behavior in larval drosophila. Learn. & Mem. 18, 639–653 (2011).

128. C Schroll, et al., Light-induced activation of distinct modulatory neurons triggers appetitive or aversive learning in Drosophila larvae. Curr. biology: CB 16, 1741–1747 (2006).

129. AS Thum, B Gerber, Connectomics and function of a memory network: the mushroom body of larval Drosophila. Curr. Opin. Neurobiol. 54, 146–154 (2019).

130. D Drix, M Schmuker, Resolving fast gas transients with metal oxide sensors. ACS sensors 6, 688–692 (2021).

131. A Spaeth, M Tebyani, D Haussler, M Teodorescu, Spiking neural state machine for gait frequency entrainment in a flexible modular robot. PloS one 15, e0240267 (2020).

